# Enhanced Loading of Paclitaxel in Cationic Liposomes by Replacement of Oleoyl with Linoleoyl Lipid Tails with Benefits in Cancer Therapeutics from *In Vitro* Studies

**DOI:** 10.1101/2020.10.12.336412

**Authors:** Yuhong Zhen, Kai K. Ewert, William S. Fisher, Victoria M. Steffes, Youli Li, Cyrus R. Safinya

## Abstract

Lipid-based carriers of the hydrophobic drug paclitaxel (PTX) are used in clinical trials as next-generation agents for cancer chemotherapy. Improving the loading capacity of these carriers requires enhanced PTX solubilization. We compared the solubility of PTX in cationic liposomes (CLs) with lipid tails containing one (oleoyl; C18:1 Δ^9^; DOTAP/DOPC) or two (linoleoyl; C18:2 Δ^9^; DLinTAP/DLinPC) *cis* double bonds with newly synthesized cationic DLinTAP (2,3-dilinoleoyloxypropyltrimethylammonium methylsufate). We used differential-interference-contrast microscopy to directly observe PTX crystal formation and generate kinetic phase diagrams representing the time-dependence of PTX solubility as a function of PTX content in the membrane. Replacing tails bearing one *cis* double bond (DO lipids) with those bearing two (DLin lipids) significantly increased PTX membrane solubility in CLs. Remarkably, 8 mol% PTX in DLinTAP/DLinPC CLs remained soluble for approximately as long as 3 mol% PTX (the membrane solubility limit which has been the focus of most previous fundamental studies and clinical trials) in DOTAP/DOPC CLs. The large increase in solubility is likely caused by enhanced molecular affinity between lipid tails and PTX upon replacement of oleoyl by linoleoyl tails, rather than by the transition in membrane structure from lipid bilayers to inverse cylindrical micelles observed in small-angle X-ray scattering. Importantly, the efficacy of PTX-loaded CLs against human prostate cancer (PC3) cells from measurements of the IC50 of PTX cytotoxicity was unaffected by changing the lipid tails, and toxicity of the CL carrier alone was negligible. Moreover, efficacy was approximately doubled against human melanoma (M21) cells for PTX-loaded DLinTAP/DLinPC over DOTAP/DOPC CLs. The findings demonstrate the potential of chemical modifications of the lipid tails to increase the PTX membrane loading well over the typically used 3 mol% while maintaining (and in some cases even increasing) the efficacy of CLs. The increased PTX solubility will aid the development of liposomal PTX carriers that require significantly less lipid to deliver a given amount of PTX, reducing side effects and costs.

## Introduction

Despite immense progress in treatment options and their effectiveness over recent decades, cancer remains a leading cause of death. Thus, there is an ongoing need for high-efficacy cancer chemotherapy with reduced side effects. Paclitaxel (PTX, Figure 1A,B)^1^ is a potent and widely used (>$1 billion/yr) cancer drug for treating ovarian, breast, lung, pancreatic, and other cancers.^1–10^ PTX inhibits mitosis by stabilizing microtubules, which are inherently dynamical *in vivo*, and subsequently activates apoptotic signaling pathways that lead to cell death.^1–3,5,11–14^

**Figure 1.**
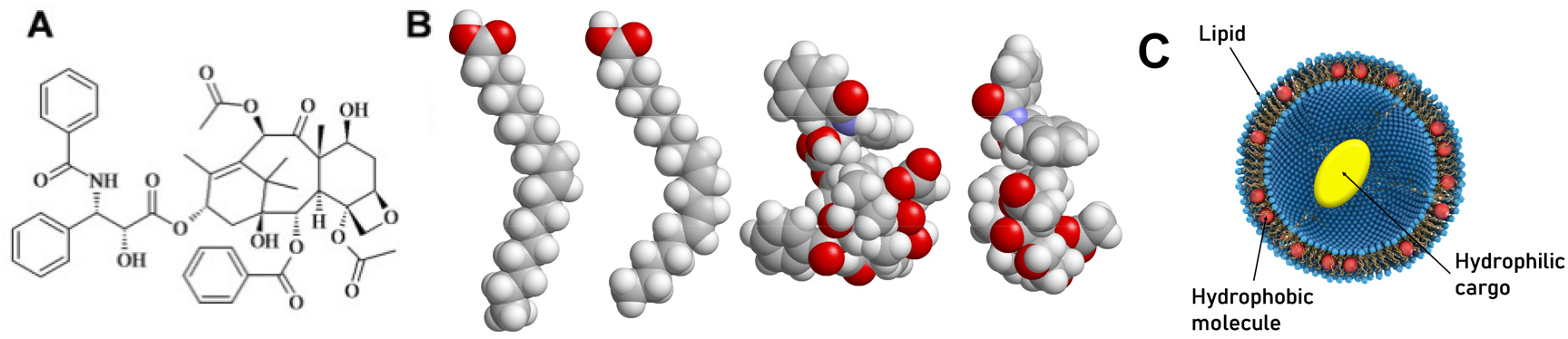
**(A)** Chemical structure of PTX. **(B)** Space filling molecular models of the ground state structure of oleic acid (C18:1) and linoleic acid (C18:2) together with two views of the structure of PTX for size comparison. **(C)** A unilamellar liposome consisting of a self-assembly of amphiphilic lipid molecules. The liposome can carry hydrophobic molecules (red spheres) within its hydrophobic bilayer and hydrophilic molecules (yellow oval) in its aqueous interior.

Because PTX is hydrophobic and poorly soluble in water, it has to be delivered by a carrier (vector).^15,16^ However, the carrier employed in the prevalent PTX formulation Taxol®,^17^ polyoxyethylated castor oil and ethanol, has been linked to severe hypersensitivity reactions requiring premedication.^18–20^ Development of more efficient and safer PTX carriers has been an ongoing challenge for decades.^21–27^ Albumin-bound PTX is an example of earlier success in carrier development and was approved by the FDA in 2005 (Abraxane®; a nontargeted nanoparticle formulation). This formulation appears to have fewer adverse reactions than Taxol and eliminates drug-carrier toxicity, but reports on whether it improves patient survival are mixed.^21,28–30^

Increasing the capacity (PTX loading) of the carrier is desirable because it means less carrier is required for a given PTX dose, reducing both cost and side effects stemming from the carrier. Furthermore, developing PTX carriers with higher efficacy, i.e., lower IC50 of PTX cytotoxicity, also reduces drug-related side effects because less PTX is required to exert its cytotoxic effect. Finally, based on its biochemical mechanism of action, PTX should be effective against most cancer cells. Therefore, development of novel, improved vectors for PTX, which, for example, may be able to deliver PTX to an expanded range of tissues, could also open treatment avenues against an expanded range of cancers. For example, Abraxane appears to be effective to treat metastatic melanoma, whereas Taxol is not.^31^

Liposomes are highly versatile and widely studied carriers of hydrophilic as well as hydrophobic drugs in therapeutic applications, in particular for cancer.^21,32–46^ Most widely-known liposomal formulations, such as Doxil and Myocet, contain the cancer drug doxorubicin in the interior of the liposome (yellow oval in Figure 1C). This is not feasible for PTX, however. PTX is much more hydrophobic (logP = 3.96) than doxorubicin (logP = 1.3), which can even be administered directly, without a solubilizing agent. In addition, the doxorubicin formulations rely on design principles that do not translate to PTX, because PTX lacks the functional groups that permit doxorubicin to be loaded via pH- or ion-gradient loading methods (forming reversibly soluble crystals within the liposomal aqueous pocket).^47^ Instead, hydrophobic drugs such as PTX are solubilized by and incorporated into the nonpolar (hydrocarbon chain) bilayer membrane of lipid-based carriers (red spheres in Figure 1C).^16,37,48–50^

Cationic liposomes (CLs; consisting of mixtures of cationic and neutral lipids) are particularly attractive as a lipid-based carrier for PTX because positively charged particles have been shown to passively target the tumor neovasculature,^38,40,43,51–56^ which has a greater negative charge than other tissues.^39,53^ Moreover, CLs are particularly attractive as a lipid-based carrier for PTX because CLs are a prevalent nonviral vector (investigated as alternatives to engineered viruses) for the delivery of therapeutic nucleic acids (NAs, e.g., plasmid DNA or siRNA; electrostatically condensed with membranes with cationic headgroups).^34,57–81^ This enables the use of CLs for combination therapies. The CL formulation EndoTAG (aka SB05), which served as a starting point for our investigations, has completed Phase II clinical trials and is currently in phase III trials.^26,52^ EndoTAG consists of CLs of the univalent cationic lipid DOTAP (2,3-dioleoyloxypropyltrimethylammonium chloride) and neutral DOPC (1,2-dioleoyl-*sn*-glycero-3-phosphatidylcholine) loaded with PTX (50:47:3 molar ratio).^37,38,40,41,82,83^

Because PTX is loaded in the bilayer of liposomes by hydrating a mixture of the lipid and PTX, the initial “loading efficiency” is 100%. However, if the amount of PTX in the membrane is larger than the membrane solubility limit, PTX precipitates out of the membrane and forms crystals over time. Once PTX has phase-separated into stable, water-insoluble crystals, the drug loses efficacy.^37,48,84–86^ Thus, it is crucial that PTX remains soluble in the membrane on timescales relevant for delivery. Relatively few studies have investigated the PTX solubility limit in different types of membranes, and not many common themes have emerged.^49,50,87–92^ Nearly all animal studies and clinical trials with liposome–PTX carriers have been conducted at 3 mol% PTX content,^37,38,41,82,83,93,94^ the first reported membrane solubility limit of PTX.^95^ Rarely, liposomal PTX formulations with higher membrane solubility have been reported, but they did not alter the structure of the lipid tails and did not provide a systematic approach to increase PTX loading.^89,96^

The PTX membrane solubility strongly depends on lipid tail structure because the location of the drug within the bilayer implies that tails exhibiting favorable local packing interactions with PTX will suppress PTX self-association, nucleation, and crystal growth. PTX is quickly expelled from membranes consisting of chain-ordered saturated lipid tails or those that have a high concentration of cholesterol.^37,49,50,87,88^ Lipids with chain-melted mono-unsaturated tails, on the other hand, are used in many of the lipid-based PTX carriers in development, such as EndoTAG®, LEP-ETU, and DHP107.^21,93^ In this work, we instead focused on tails with multiple *cis* double bonds because these increase chain disorder and modify chain flexibility. We hypothesized that this increased chain disorder and altered flexibility would affect molecular affinity to PTX compared to tails with one *cis* double bond. Currently there are only a few instances of commercial therapeutics containing lipids with poly-unsaturated fatty acid tails.^32,97^ A notable exception is DLin-MC3-DMA (Figure 2), the cationic lipid component of patisiran (Onpattro®),^98^ which became the first FDA-approved siRNA therapeutic in 2018. We note, however, that the siRNA component interacts with the lipid’s headgroup in the case of patisiran and thus, unlike in our system, the active ingredient has no direct interactions with the polyunsaturated tails. (As an aside, poly-unsaturated fatty acids have been studied as therapeutic entities in and of themselves for their anti-oxidant properties.^99,100^)

**Figure 2.**
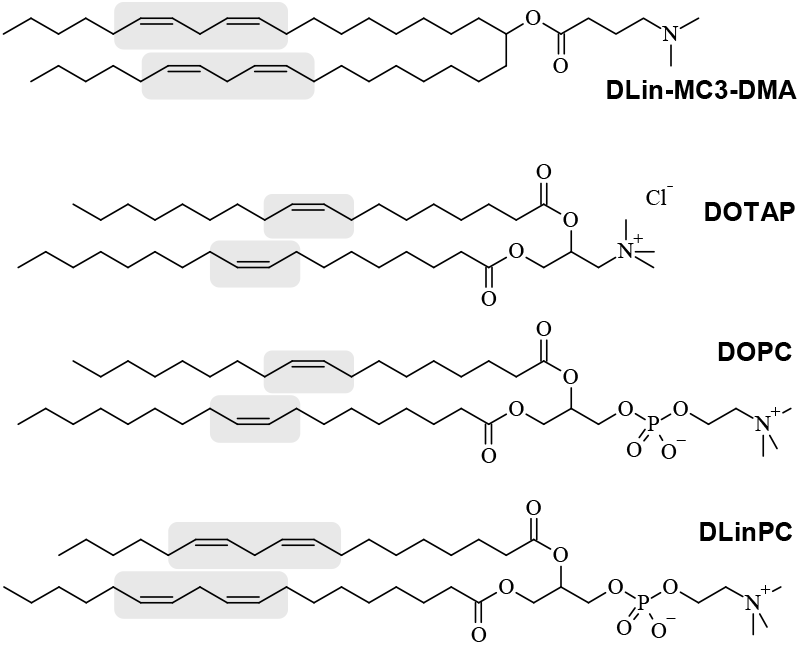
Chemical Structures of DLin-MC3-DMA (the cationic lipid used in patisiran), DOTAP, DOPC (the lipids used in the Endo-TAG formulation of PTX) and DLinPC. The *cis* double bonds in the lipid tails are highlighted.

We pursued the development of CL carriers with a tail structure that improves solubility of PTX in their hydrophobic membrane. Such vectors require less lipid to deliver a given amount of PTX, reducing costs and side effects. Carriers with high solubility have also shown increased efficacy,^48^ allowing administration of lower total doses of PTX. We obtained promising initial results of increased PTX membrane solubility using CLs prepared from DOTAP (oleoyl (C18:1) tails; see Figure 2) and DLinPC (linoleoyl (C18:2) tails; see Figure 2) (at a molar ratio of DOTAP/DLinPC/PTX=30/70–x/x).^101^ Encouraged by these results, we synthesized the univalent cationic lipid DLinTAP from linoleic acid and used it to prepare PTX-loaded CLs containing lipids with exclusively C18:2 tails.

We used differential-interference-contrast (DIC) microscopy to directly observe PTX crystal formation and generate kinetic phase diagrams, characterizing the time-dependence of PTX solubility as a function of PTX content for CLs with lipid tails containing either one (DOTAP/DOPC) or two (DLinTAP/DLinPC) *cis* double bonds. Using cell viability measurements, we then compared the efficacy of PTX-loaded DLinTAP/DLinPC and DOTAP/DOPC CLs in PC3 (prostate) and M21 (melanoma) human cancer cell lines and determined the IC50 for PTX cytotoxicity of these vectors in the cell lines.

Replacing tails bearing one *cis* double bond (DO lipids) with those bearing two (DLin lipids) significantly increased PTX membrane solubility in CLs. Remarkably, 8 mol% PTX in DLinTAP/DLinPC CLs remained soluble for approximately as long as 3 mol% PTX in DOTAP/DOPC CLs. At the same time, the IC50 of PTX cytotoxicity against PC3 cells was unaffected by changing the lipid tails, and toxicity of the lipid carrier was negligible. In M21 cells, efficacy was not just maintained but approximately doubled for PTX-loaded DLinTAP/DLinPC CLs over DOTAP/DOPC CLs.

Small-angle X-ray scattering (SAXS) allowed determination of the self-assembled nanostructures of PTX-containing CLs, mixed with DNA as a condensing agent to improve the SAXS signal, at varying amounts of DLinPC and DOTAP. The structures begin to transition from lamellar (L_α_^C^) to inverse hexagonal (H_II_^C^) as the content of DLinPC is increased to 70 mol% DLinPC and beyond. However, kinetic phase diagram studies show that PTX membrane solubility decreases in membranes with dioleoyl tails upon transition from the lamellar to the H_II_ phase by replacing DOPC with DOPE in DOTAP-containing CLs. These results suggest that the significantly improved PTX membrane solubility in DLinTAP/DLinPC CLs is caused by enhanced affinity of PTX to linoleoyl compared to oleoyl tails, rather than by the structural transition from lipid bilayers to inverse cylindrical micelles.

Taken together, our findings show that CLs with suitable tail structure can increase the PTX membrane loading from the typically used 3 mol% to approximately 8 mol%, and that the efficacy of those CLs is as high or twice as high as the Endo-TAG formulation for PC3 cells and M21 cells, respectively. This will aid the development of liposomal PTX carriers that use less lipid, reducing side effects and costs.

## Materials and Methods

### Materials

Stock solutions of DOPC, DOTAP, and DLinPC in chloroform were purchased from Avanti Polar Lipids. PTX was purchased from Acros Organics and dissolved in chloroform to 10.0 mM concentration. Calf thymus DNA was purchased from Thermo Scientific. DLinTAP was synthesized as described in the supplementary material.

### Cell Culture

The human prostate cancer cell line PC3 (ATCC number: CRL-1435) and human melanoma cell line (M21) were a gift from the Ruoslahti Lab (Sanford Burnham Prebys Medical Discovery Institute). M21 cells are a subclone of the human melanoma line UCLA-SO-M21 derived in the lab of R. Reisfeld (Scripps Institute, La Jolla) and originally provided by D. L. Morton (UCLA). Cells were cultured in DMEM (Invitrogen) supplemented with 10% fetal bovine serum (Gibco) and 1% penicillin/streptomycin (Invitrogen). Cells were cultured at 37°C in a humidified atmosphere with 5% CO_2_ and split at a 1:5 ratio after reaching ≥80% confluency (every 48-72 hours) during maintenance.

### Liposome Preparation

Suspensions of sonicated and unsonicated liposomes at a total molar concentration (lipid + PTX) of 1 mM for cell viability experiments, 5 mM for DIC microscopy, and 30 mM for small-angle X-ray scattering measurements were prepared as described previously.^48^

### DIC Microscopy (PTX Solubility and Kinetic Phase Diagrams)

Samples were prepared and assayed as described previously,^48^ with the following modifications: the sample solutions were stored at room temperature for the duration of the experiment and imaged at 20 or 40 × magnification every 2 h until 24 h, every 12 h until 48 h, and daily thereafter until PTX crystals were observed. The kinetic phase diagrams report the time at which PTX crystals were observed in 2 of 3 independently prepared samples at each PTX content.

### Small-Angle X-Ray Scattering

Samples for X-ray scattering were prepared by combining and vortexing 50 μL of a 30 mM aqueous liposome suspension with DNA solution (3.5 mg/mL in water) at a cationic lipid to DNA charge ratio of 1 in a 500 μL centrifuge tube. Following centrifugation in a table-top centrifuge at 5,000 rpm, the resulting pellets were transferred to quartz capillaries (Hilgenberg) with the help of excess supernatant. The capillaries were then centrifuged in a capillary rotor in a Universal 320R centrifuge (Hettich) at 10,000 *g* and 25 °C for 30 min. After centrifugation, the capillaries were sealed with a fast-curing epoxy glue.

SAXS measurements were carried out at the Stanford Synchrotron Radiation Laboratory, beamline 4-2, at 9 keV (λ=1.3776 Å) with an Si(111) monochromator. Scattering data was measured by a 2D area detector (MarUSA) with a sample to detector distance of ≈3.5 m (calibrated with silver behenate). The X-ray beam size on the sample was 150 μm in the vertical and 200 μm in the horizontal directions. Scattering data is reported as azimuthally averaged scattering intensity in *q*-space.

### Cell Viability Assays

Cells were plated at a density of 5,000 cells/well in 96-well plates and incubated overnight. Suspensions of sonicated liposomes were diluted in DMEM to reach the desired PTX concentration. The culture medium was removed from the wells by aspiration with a pipette, taking care not to aspirate any cells, and a total of 100 μL of the liposome suspension in DMEM was added to each well. Cells were incubated for 24 h before the liposome suspension was removed by manual pipetting and replaced with cell culture medium. After incubation for 48 h, the cell viability was measured using the CellTiter 96® AQueous-One Soution Cell Proliferation Assay (Promega). The assay solution was diluted 6-fold with DMEM, and a total of 120 μL of this solution was added to each well. After 1 h of incubation, absorbance was measured at 490 nm using a plate reader (Tecan M220). Each data point shown comprises four identically treated wells and is normalized to the absorbance obtained for untreated cells.

Liposomes with different PTX contents were added to cells such that the resulting PTX concentration per well was identical for each data point, independent of the formulation. Because of this, the lipid concentration at each data point varies between formulations with different PTX contents.^48^ To determine the IC50 values, we used the solver add-in for Microsoft Excel to perform a nonlinear least squares fit of the cell viability data to the equation y=A+B/(1+[x/C]^D^). Here, y is the measured (normalized) cell viability, x is the total PTX concentration, A is the minimum cell viability, (A+B) is the maximum cell viability (B is the range of y), C is the IC50 (the concentration of PTX at which cell viability is halfway between the maximum and minimum values, i.e. where y=A+B/2), and D is the “slope factor” of the curve (indicating how steeply the viability declines). The minimum and range of cell viabilities (A and B) was given by the data, while C and D were used as fitting parameters.

## Results and Discussion

### Lipid Synthesis

**Scheme 1.**
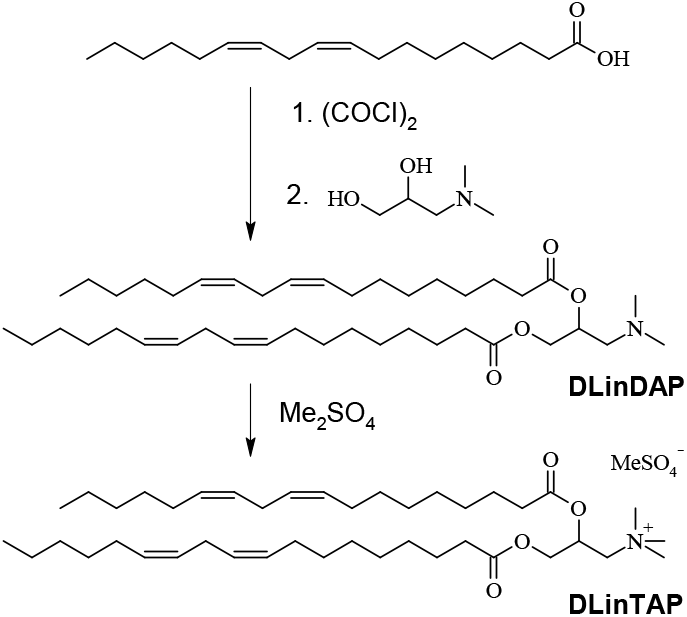
Synthesis of DLinTAP. The full details of the synthesis are reported in the supplementary material.

To be able to prepare CLs with tails that exclusively bear two *cis* double bonds (derived from linoleic acid, C18:2 Δ^9^), we synthesized DLinTAP as shown in Scheme 1. The full details of the synthesis, which used a route analogous to that reported for DOTAP,^102^ are reported in the supplementary material.

### PTX Membrane Solubility

Upon hydration of films prepared from a mixture of cationic lipid (DLinTAP or DOTAP), neutral lipid (DLinPC or DOPC) and PTX, PTX-loaded CLs formed spontaneously. These CLs were studied directly (unsonicated CLs; uni- and multi-lamellar with a broad distribution of larger sizes and an average diameter of ≈800 nm) or after sonication (small unilamellar CLs with diameter < 200 nm). As mentioned in the Introduction, the initial “loading efficiency” is 100%, but, depending on the composition, PTX may phase separate and crystallize over time, decreasing the PTX loading of the CLs and reducing the amount of PTX that is effective against cancer cells.

#### DIC Microscopy

To compare the solubility of PTX in CLs prepared from DLin- and DO-lipids, we used differential interference contrast (DIC) microscopy.^48^ Starting 2 h after sample hydration, samples were observed at regular time intervals to check for PTX crystals as evidence of phase separation. Figure 3 shows DIC micrographs illustrating the variety of size and shape in the observed crystals (for unsonicated samples). In addition to characteristic needle-shaped PTX crystals (Figure 3A,D,E), we commonly observed star and double-fan shapes (Figure 3B, D and Figure 3C, respectively). As PTX content increased, the number of crystals increased and their size decreased (compare Figure 3B to Figure 3E). Aggregates of PTX crystals were common at contents above 6 mol%, including the feather-like crystals observed at 9 mol% (Figure 3F). The aspect ratio of PTX crystals from DLinTAP/DLinPC CLs was typically much smaller than that of crystals formed from DOTAP-containing CLs (compare Figure 3A,D to Figure 3B,C,E). The overall smaller size and larger number of PTX crystals from DLinTAP/DLinPC CLs suggests that there are fewer nucleation and growth sites in DOTAP-based samples.

**Figure 3.**
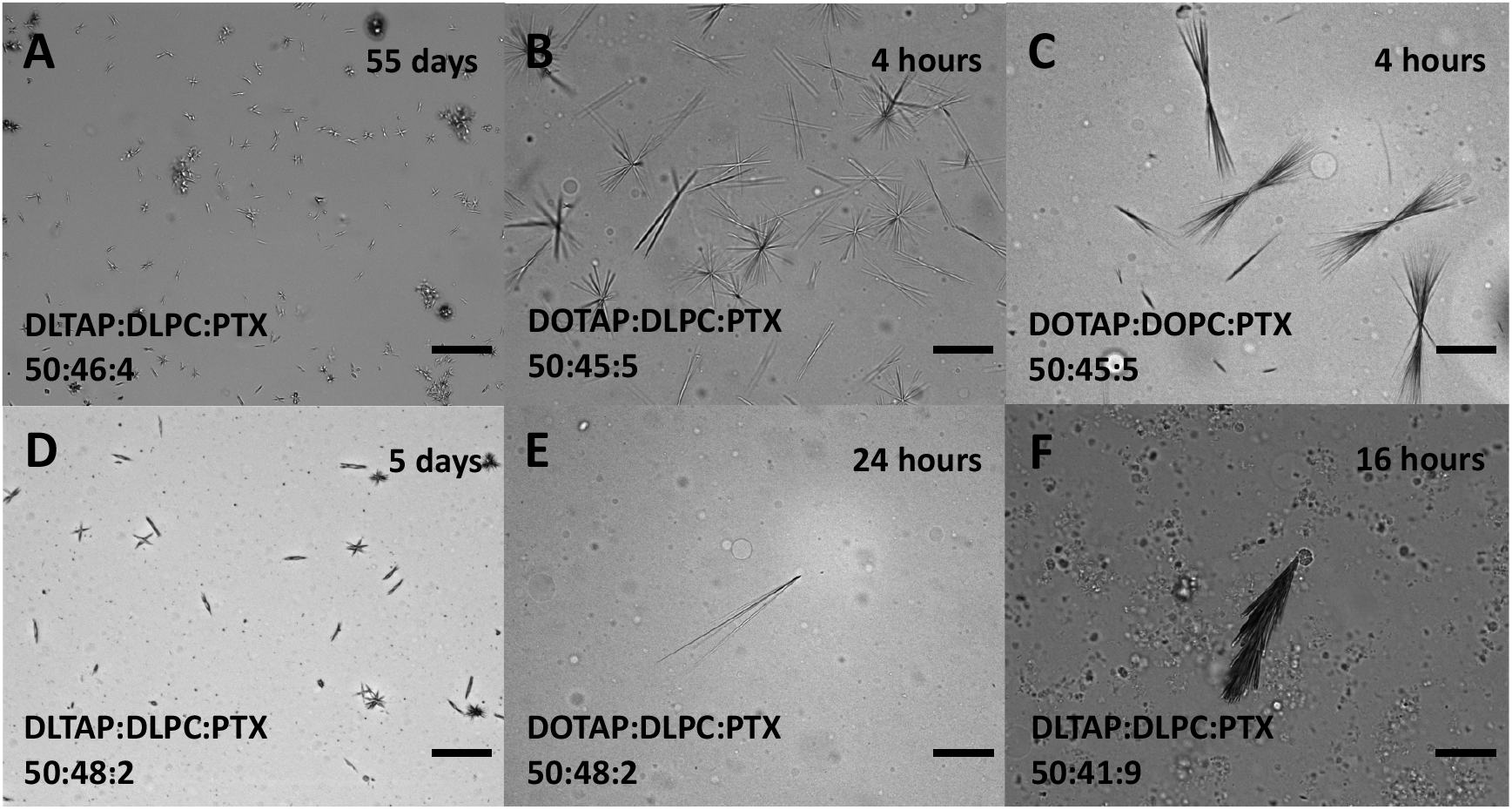
Selected DIC microscopy images of liposomes and phase separated PTX, illustrating the size and shape variety of observed PTX crystals as discussed in the text. The samples were unsonicated PTX-loaded CLs of the molar compositions indicated on the images. Images were taken after hydration for the time noted. Scale bars: 50 μm.

#### Kinetic Phase Diagrams of PTX-Loaded CLs

The time-dependent PTX solubility data as a function of PTX content (kinetic phase diagrams) of sonicated and unsonicated PTX-loaded DLinTAP/DLinPC CLs (DLinTAP/DLinPC/PTX mole ratio=50:50–x_PTX_:x_PTX_), as mapped out by DIC microscopy, are shown in Figure 4A and 4B, respectively. Blue color indicates that PTX remained solubilized in the CL membranes, i.e., no crystals were observed. Red color indicates time points at which PTX crystals were observed. (Time points after the first observation of crystals were marked with red color even if no further samples were assessed, since crystallization from the membrane is irreversible.) For example, in the sonicated sample containing 10 mol% PTX (DLinTAP/DLinPC/PTX mole ratio=50:40:10), crystals were first observed at the 6 h time point, meaning that PTX crystallized between 4 and 6 h after hydration (Figure 4A). The line separating the blue and red regions of the kinetic phase diagram, which marks the onset of crystal formation, is termed the PTX membrane solubility boundary. For comparison, the black lines in Figure 4A and 4B indicate the membrane solubility boundary for PTX-loaded DOTAP/DOPC CLs (where x_PTX_=3 is the Endo-TAG composition).

**Figure 4.**
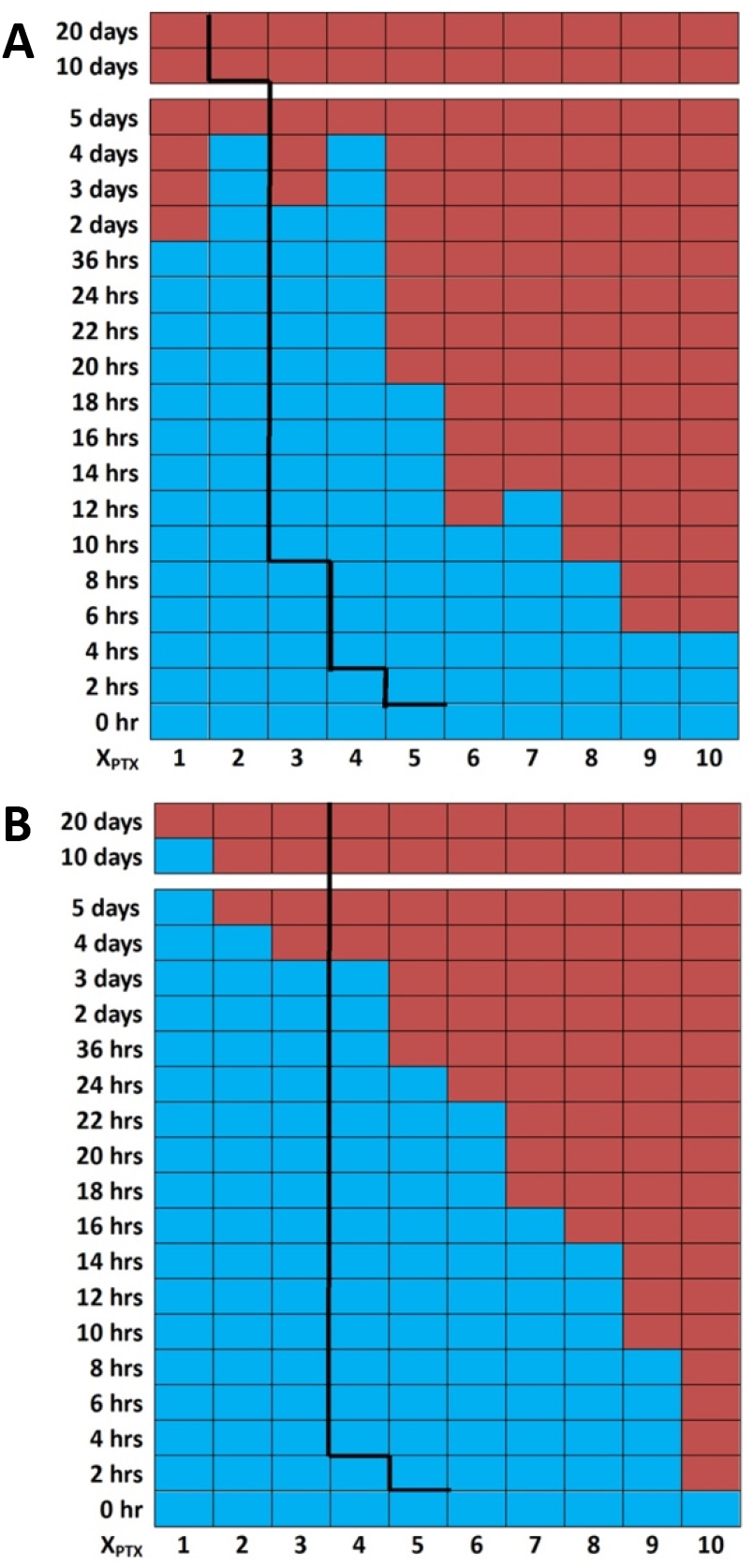
Kinetic phase diagrams of PTX solubility in PTX-loaded CLs with DLin (di-linoleoyl, C18:2) lipid tails (DLinTAP/DLinPC/PTX mole ratio=50:50–x_PTX_:x_PTX_). DIC microscopy (Figure 3) was used to detect PTX crystallization at the indicated times after hydration. The blue color indicates absence of PTX crystals (i.e., PTX remained soluble in the membranes), while the red color indicates presence of PTX crystals. The boundary separating the blue region from the red region is termed the PTX membrane solubility boundary. As a reference to facilitate comparison, the black lines show the solubility boundary for PTX-loaded CLs with monounsaturated DO (di-oleoyl, 18:1) tails (DOTAP/DOPC/PTX molar ratio=50:50-x_PTX_:x_PTX_). The change in tail structure strongly affects PTX membrane solubility. **(A)** Kinetic phase diagram for sonicated PTX-loaded DLinTAP/DLinPC CLs. **(B)** Kinetic phase diagram for unsonicated PTX-loaded DLinTAP/DLinPC CLs.

It is immediately obvious from inspection of the kinetic phase diagrams that the switch from DO- to DLin-tails dramatically improves the solubility of PTX in the CL membrane. For sonicated CLs (Figure 4A), for example, a PTX content of 8 mol% remained solubilized in DLinTAP/DLinPC CLs for as long as a content of 3 mol% PTX did in DOTAP/DOPC CLs. The duration of PTX solubility in unsonicated CLs at most PTX contents is longer than in sonicated CLs, independent of the tail structure. However, CLs with DO tails show a solubility threshold at 3 mol% PTX content, meaning that PTX had crystallized from CLs with PTX content of 4 mol% at 4 h after hydration for both sonicated and unsonicated samples. (This is consistent with the reported PTX membrane solubility limit of 3 mol% used for most liposomal PTX vectors.) In contrast, the kinetic phase diagrams show a much more gradual decrease in the time for which PTX stayed solubilized with increasing PTX content for CLs with DLin tails. This is especially noticeable for unsonicated liposomes (Figure 4B), where PTX remained soluble in DLinTAP/DLinPC CLs for over 3 days at 4 mol% and for over 22 h (but less than 24 h) at 6 mol%. Even at 9 mol%, PTX only crystallized between 8 and 10 h after hydration. What is impressive is that for sonicated CLs (similar in size and size distribution to preparations used in clinical trials) at 8 mol% PTX, crystals appeared on average only between 8 and 10 h after hydration. This relatively long time of PTX solubility should be sufficient for most *in vivo* applications.

Only at low loadings (PTX content <2 mol% and <3 mol% in sonicated and unsonicated samples, respectively) and long incubation times was PTX more soluble in DOTAP/DOPC CLs than in DLinTAP/DLinPC CLs. This may be due to oxidation of the DLin tails, because we took no special precautions to exclude oxygen from the small sample volumes during the repeated withdrawing of aliquots for DIC microscopy over time.

### Small-Angle X-Ray Scattering

To investigate the effect of tail saturation on the structure of CL membranes, we used synchrotron small-angle X-ray scattering (SAXS) to determine the structure of CLs prepared from mixtures of DOTAP with either DOPC or DLinPC. To enhance the signal-to-noise ratio, we condensed the CLs with DNA. This has been shown to result in CL–DNA complexes where the equilibrium self-assembled structure of the membrane within the complex is determined by the spontaneous curvature (*C*_0_) of the lipid self-assembly.^34,103-105^ The spontaneous curvature is, in turn, determined by the average shape of the lipid molecules.^106^

DOPC/DOTAP CLs mixed with DNA form the lamellar (L_α_^C^) phase because both DOPC and DOTAP have a cylindrical shape with *C*_0_≈0 (see Figure 6).^103^ We expected that DLinPC could be capable of forming the inverse hexagonal (H_II_) nonbilayer structure due to the increase in unsaturation from one to two *cis* double bonds in the lipid tails. The two *ci*s double bonds in DLinPC induce kinks in the lipid tails that can not readily be offset by *gauche* conformations in the single bonds, leading to the tails taking up a bigger lateral area compared to the headgroup area (Figure 6). This results in an inverted-cone molecular shape as depicted in Figure 6, corresponding to negative spontaneous curvature (*C*_0_ < 0). A previous study on soy PC (a lipid mixture largely composed of DLinPC) with x-ray scattering and cryogenic TEM supports this hypothesis.^107^

Figure 5 shows the azimuthally-averaged scattering profiles as a function of the reciprocal lattice vector *q* for CL–DNA complexes with membrane compositions of 70 or 80 mol% neutral lipid, 2 mol% PTX, and the remainder DOTAP. Peak assignments are shown on the plots as follows: L_α_^C^ (00L: 001, 002, 003, 004, 005, 006), H_II_^C^ (HK: 10, 20, 21, 30, 31, 40), DNA–DNA interaxial spacing in the L_α_^C^ phase (DNA). Additionally, three peaks that can be assigned to crystallized PTX are visible in the sample with 70 mol% DLinPC (*q*_P1_ = 0.291 Å^−1^, *q*_P2_ = 0.374 Å^−1^, and *q*_P3_ = 0.496 Å^−1^), consistent with previous studies.^48^ The L_α_^C^ phase consists of alternating lipid bilayers and DNA^103^ and the H_II_^C^ phase consists of a hexagonal arrangement of inverse cylindrical micelles with DNA inserted in their lumen.^104^

**Figure 5.**
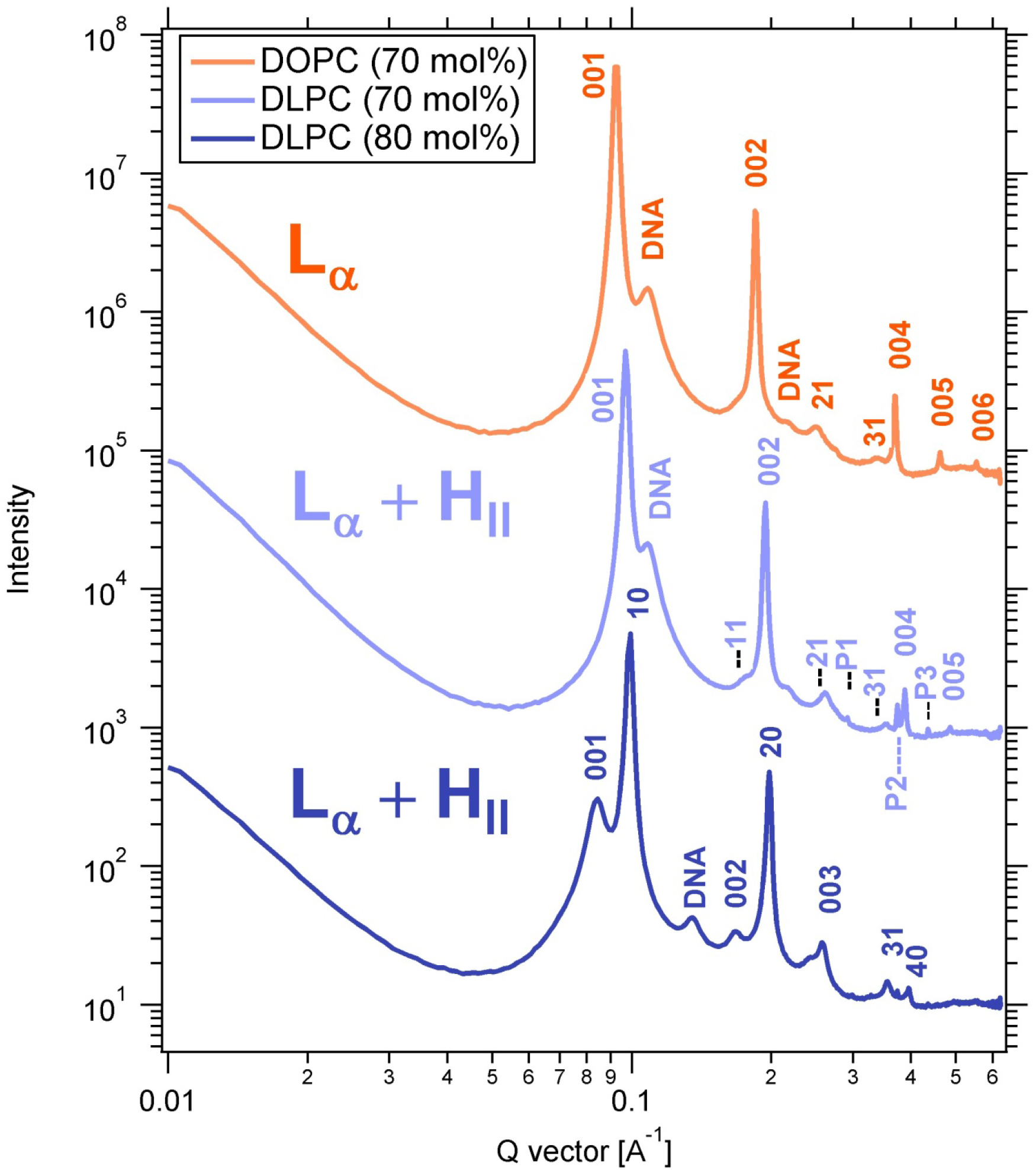
X-ray scattering profiles CL–DNA complexes prepared from PTX-loaded CLs with high contents of DOPC or DLinPC, revealing their self-assembled structures. Peak assignments shown are for L_α_^C^ (00L), H_II_^C^ (HK), DNA–DNA interaxial spacing in the L_α_^C^ phase (DNA), and crystallized PTX (P1, P2, P3). The CLs were composed of 70 or 80 mol% neutral lipid (DOPC or DLinPC), 2 mol% PTX, and the remainder DOTAP and were complexed with calf thymus DNA at a 1:1 charge ratio.

The X-ray scattering profile from CL–DNA complexes with 70 mol% DOPC shows peaks characteristic of the lamellar L_α_^C^ phase (Figure 5, top profile). Based on this result and previous reports,^108^ we expect that CL–DNA complexes with 80 mol% DOPC in the membrane would also be L_α_^C^ phase. In contrast, SAXS showed that CL–DNA complexes with 70 and 80 mol% of DLinPC in the membrane were in a 2-phase regime, consisting of a mixture of the lamellar (L_α_^C^) and inverted hexagonal (H_α_^C^) phases. This is expected because the cylindrical DOTAP with *C*_0_≈0 prefers the L_α_^C^ phase, whereas the inverse-cone-shaped DLinPC with *C*_0_<0 prefers the H_II_^C^ phase. The peak intensities (Figure 5) reveal that the L_α_^C^ phase dominates at 70 mol% DLinPC, but at increased DLinPC content of 80 mol%, the H_II_^C^ phase is dominant.

Interestingly, at 70 mol% neutral lipid (Figure 5, top two profiles), the *q*_001_ peak of the L_α_^C^ phase in the sample containing DLinPC is observed at higher *q* (*q*_001_=0.0968 Å^−1^) than the peak of the sample with DOPC (*q*_001_=0.0931 Å^−1^). The interlayer spacing *d*_*lamellar*_=2π/*q*_001_ correlates to the membrane thickness. Thus, the shift in *q*_001_ indicates that the interlayer spacing drops from 67.5 Å to 64.9 Å when DOPC is replaced with DLinPC. DLinPC membranes are therefore thinner than DOPC membranes, which is consistent with the expectation that the added double bond forces the tails to take up more lateral width, thereby thinning the membrane (as shown in Figure 6).

**Figure 6.**
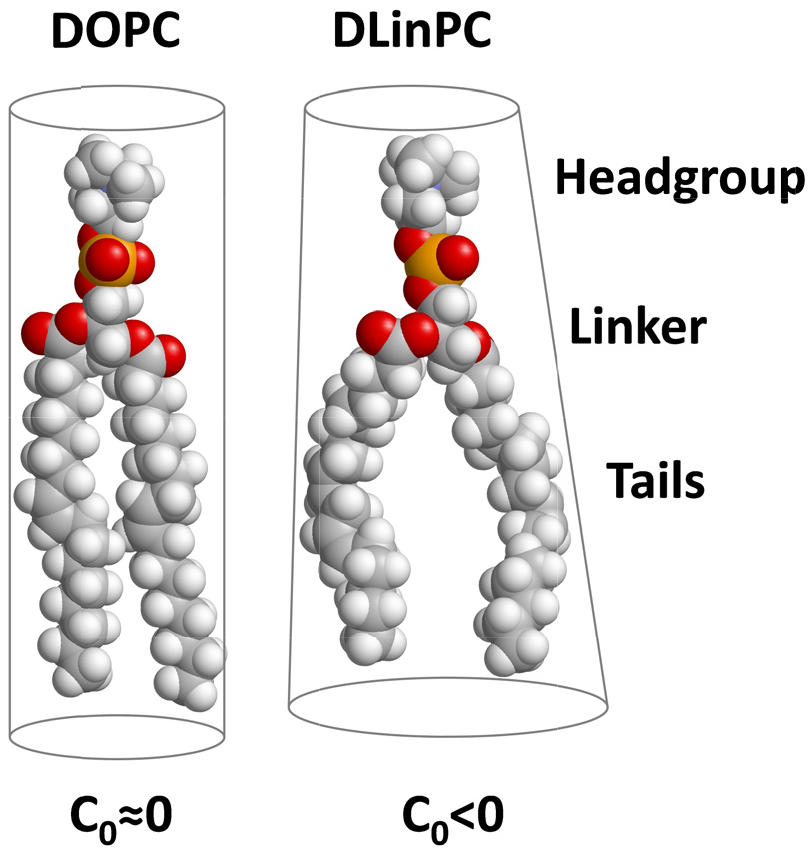
Effect of increased tail unsaturation on lipid molecular shape. The additional *cis* double bond in the linoleoyl tails induces a kink in the tails that is not readily offset by gauche conformations of single bonds in the chain. The linoleoyl tails therefore take up a greater lateral area. This changes the molecular shape, resulting in a different spontaneous curvature (*C*_0_<0) and thinning f the bilayer.

The results of the SAXS studies could indicate that the significantly improved PTX membrane solubility upon replacing oleoyl tails in CLs with linoleoyl is due to the structural transition from lipid bilayers to inverse cylindrical micelles. However, PTX solubility is actually much lower in membranes that form the H_II_^C^ instead of the L_α_^C^ phase if they consist only of oleoyl tails. This is evident from Figure S1 in the supplementary material, which shows the kinetic phase diagram for PTX solubility in CLs containing 70 mol% DOPE, DOTAP and PTX. DNA complexes of these membranes form the H_II_^C^ phase (see Figure S2 in the supplementary material) because the headgroup of DOPE (phosphoethanolamine) is smaller than that of DOPC(phosphocholine). Therefore, improved solubility of PTX in DLinTAP/DLinPC membranes is not due to their preference for the HIIC phase but rather the different interactions of DLin tails with PTX.

### Cytotoxicity

Our kinetic phase diagram data suggests that CLs composed of DLinTAP and DLinPC can stably solubilize up to nearly 3-fold more PTX in their membrane than those composed of DOTAP and DOPC (Figure 4). This enhanced loading capacity will only provide significant benefits if the amount of PTX required to achieve cytotoxic efficacy against cancer cells does not change. To compare the ability of DOTAP/DOPC and DLinTAP/DLinPC PTX carriers to induce cancer cell death (i.e., their efficacy), we measured their IC50 for PTX cytotoxicity in cell viability experiments for two human cancer cell lines. The IC50 is defined as the concentration of PTX required to elicit half the maximum cytotoxic effect. These cytotoxicity experiments were conducted using sonicated CLs.

Figure 7 shows results from cell viability experiments using CLs composed of DLinTAP/DLinPC and loaded with PTX to target prostate cancer metastasis (PC3) cells. For these measurements, we varied the PTX content of the CLs to span the range of PTX membrane solubility observed in the kinetic phase diagrams. Specifically, the PTX content ranged from 2 mol%, which is stably solubilized in both DOTAP/DOPC and DLinTAP/DLinPC CLs, up to 9 mol%, which rapidly precipitates from DOTAP/DOPC CLs and is only moderately stable in DLinTAP/DLinPC CLs. The viability of PC3 cells (normalized to untreated cells) rapidly decreases with increasing PTX concentration delivered by DLinTAP/DLinPC CLs (Figure 7A), with most of the decrease occurring between 5 and 20 nM PTX.

**Figure 7.**
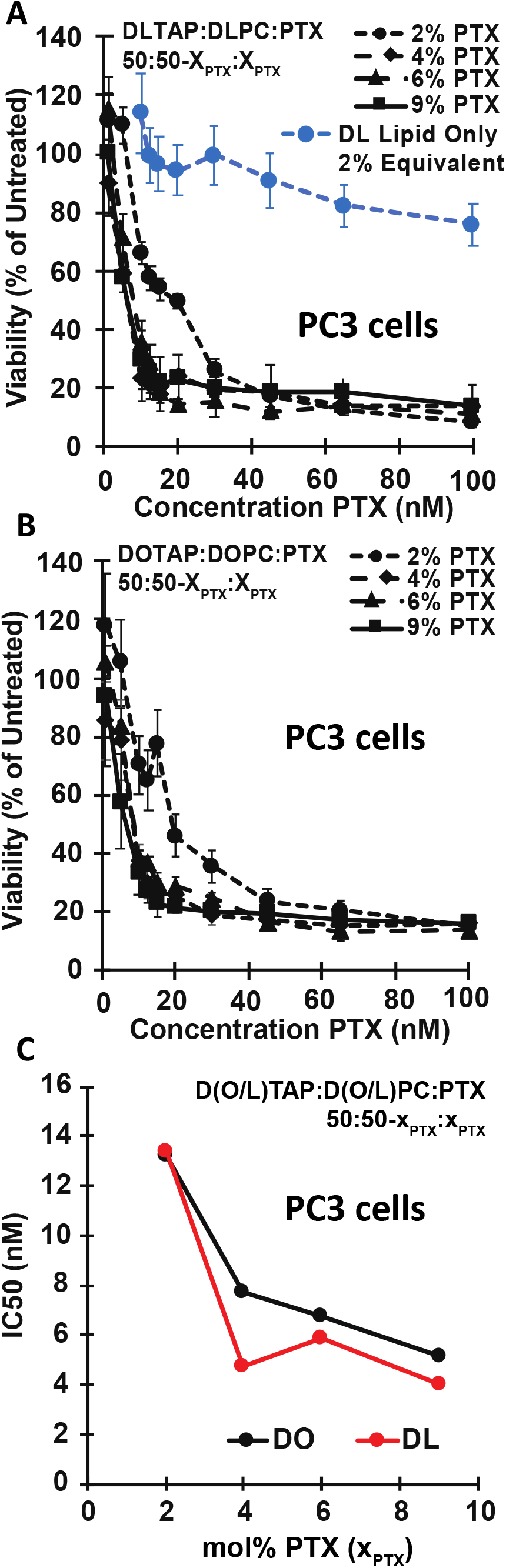
Cytotoxicity of PTX-loaded CLs against PC3 cells. **A)** Viability of PC3 cells (relative to untreated cells) as a function of PTX concentration for cells treated with CLs of a molar composition of 50/50– x_PTX_/x_PTX_, DLinTAP/DLinPC/PTX, with x_PTX_=2, 4, 6, or 9 (black lines). Cell viability rapidly decreased with PTX concentration to a plateau. As a control, DLinTAP/DLinPC CLs without PTX were added to PC3 cells in amounts that match the lipid content of CLs with x_PTX_=2 (which have the highest lipid content) at the PTX concentrations tested (blue line). This CL-only control showed no appreciable cytotoxicity (≥90% viability) up to the lipid equivalent of 65 nM PTX delivered with a 2 mol% PTX formulation, demonstrating that PTX and not lipid drove cytotoxicity at the IC50 of every formulation. **B)** Viability of PC3 cells (relative to untreated cells) as a function of PTX concentration for cells treated with control CLs of a molar composition of 50/50–x_PTX_/x_PTX_, DOTAP/DOPC/PTX, with x_PTX_=2, 4, 6, or 9 (black lines). Cell viability rapidly decreased with PTX concentration to a plateau. Previous work has shown that CLs alone do not contribute to cytotoxicity at the employed concentrations.^48^ **C)** Plot of IC50 values for PTX cytotoxicity against PC3 cells for the CL formulations based on lipids with linoleoyl (“DLin”, red line) and oleoyl (“DO”, black line) tails. Each IC50 was determined by fitting the corresponding cell viability curve (from parts A and B) as described in the Methods section. The IC50 decreases (efficacy increases) with increasing PTX content in both DOTAP/DOPC/PTX and DLinTPA/DLinPC/PTX formulations. The efficacy of DLinTAP/DLinPC/PTX CLs is higher than that of DOTAP/DOPC/PTX CLs at all PTX contents except 2 mol%, but difference is not significant. This shows that replacing oleoyl with linoleoyl tails maintains cytotoxic efficacy against PC3 cells while increasing PTX membrane solubility (Figure 4).

To control for lipid toxicity, we also measured the cell viability for increasing concentrations of DLinTAP/DLinPC CLs without any PTX (Figure 7A, blue curve). For comparison with PTX-loaded CLs, we incubated the cells with the amount of DLinTAP/DLinPC CLs required to deliver PTX at 2 mol% PTX content at each PTX concentration (because the formulation at 2 mol% PTX has the highest CL/PTX ratio). Lipid toxicity (>10% drop in cell viability) was observed only at lipid concentrations equivalent to or greater than those required to deliver 65 nM PTX at 2 mol% PTX content. Because the IC50 for PTX-loaded DLinTAP/DLinPC CLs at 2 mol% PTX was 13.4 nM, more than four times lower than the PTX concentration where lipid toxicity was observed with this formulation, it is unlikely that lipid toxicity contributed to its cytotoxic efficacy. The same is true for all other formulations, considering that they delivered more PTX per lipid and had lower IC50 values than the CLs with 2 mol% PTX content.

To compare the cytotoxic efficacy of DLinTAP/DLinPC CLs to that of CL-based PTX carriers currently in clinical use, we measured the cytotoxicity of PTX-loaded DOTAP/DOPC CLs (Figure 7B). These CLs also decreased the viability of PC3 cells (normalized to untreated cells) as a function of delivered PTX concentration, with a rapid drop of cell viability between 5 and 20 nM PTX. Steffes et al. previously demonstrated^48^ that DOTAP/DOPC CLs without PTX are not cytotoxic at any lipid concentration used in this study and therefore unlikely to affect the cytotoxicity of the PTX-loaded CLs.

To facilitate comparison of the DLinTAP/DLinPC and DOTAP/DOPC formulations, Figure 7C plots their IC50 for cytotoxicity of PTX against PC3 cells as a function of their PTX content. The IC50 values of DLinTAP/DLinPC and DOTAP/DOPC CLs at each PTX content are very similar, demonstrating that cytotoxic efficacy is unaffected by the change in tail structure. The efficacy against PC3 cells increased (i.e., IC50 decreased) with increasing PTX content in formulations with both DO- and DLin-lipids, while previous studies had found an increase in the IC50 with PTX content for DO-lipid-based CLs.^48^ A possible explanation for this is that, even at high PTX content, the PTX remained solubilized in the membranes long enough to exert its cytotoxic effects, allowing successful PTX delivery by metastable CLs. If such PTX-loaded CLs are not used immediately after preparation, however, PTX crystallizes and their efficacy drops.^48^

We also investigated the efficacy of PTX-loaded DLinTAP/DLinPC CLs against a metastatic melanoma (M21) cell line, using the same range of CL formulations and PTX contents as for the PC3 cell line (Figure 8A). Compared to PC3 cells, the viability of M21 cells decreased more gradually, with most of the drop in viability occurring between 10 and 65 nM PTX, depending on the specific formulation. This is consistent with prior investigations using PTX-loaded DOTAP/DOPC CLs.^48^

**Figure 8.**
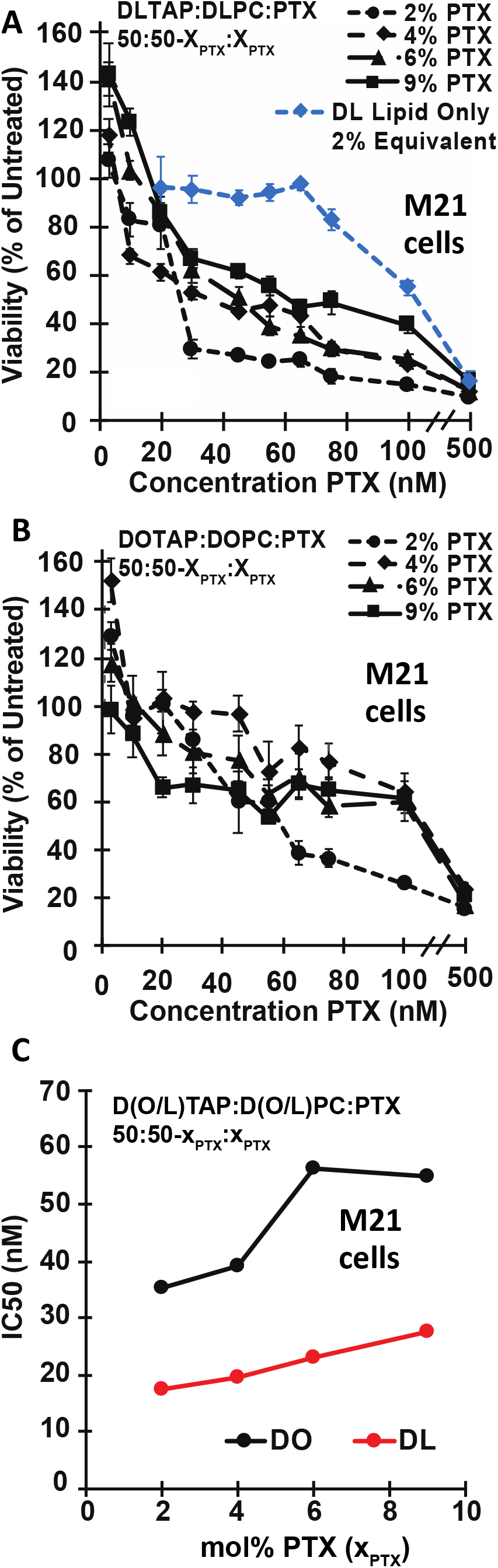
Cytotoxicity of PTX-loaded CLs against M21 cells. **A)** Viability of M21 cells (relative to untreated cells) as a function of PTX concentration for cells treated with CLs of a molar composition of 50/50– x_PTX_/x_PTX_, DLinTAP/DLinPC/PTX, with x_PTX_=2, 4, 6, or 9 (black lines). Cell viability decreased with PTX concentration to a similar level as observed for PC3 cells, but did so more gradually. As a control, DLinTAP/DLinPC CLs without PTX were added to M21 cells in amounts that match the lipid content of CLs with x_PTX_=2 (which have the highest lipid content) at the PTX concentrations tested (blue line). This CL-only control showed no appreciable cytotoxicity (≥90% viability) up to the lipid equivalent of 75 nM PTX delivered with a 2 mol% PTX formulation, suggesting that PTX and not lipid drove cytotoxicity at the IC50 of every formulation because CL cytotoxicity affects cell viability only at concentrations at least four times higher than the IC50 (see part C). **B)** Viability of M21 cells (relative to untreated cells) as a function of PTX concentration for cells treated with control CLs of a molar composition of 50/50–x_PTX_/x_PTX_, DOTAP/DOPC/PTX, with x_PTX_=2, 4, 6, or 9 (black lines). The decrease in cell viability with PTX concentration is less steep than in part A, suggesting a lower efficacy. Previous work has shown that CLs alone do not contribute to cytotoxicity at the employed concentrations.^48^ **C)** Plot of IC50 values for PTX cytotoxicity against M21 cells for the CL formulations based on lipids with linoleoyl (“DLin”, red line) and oleoyl (“DO”, black line) tails. Each IC50 was determined by fitting the corresponding cell viability curve (from parts A and B) as described in the Methods section. Importantly, the efficacy of the DLinTAP/DLinPC/PTX formulations was about two-fold higher (their IC50 values were two-fold lower) than that of the corresponding DOTAP/DOPC/PTX formulations. This effect amplifies the benefits that replacing oleoyl with linoleoyl tails cells brings by to increasing PTX membrane solubility (Figure 4). In contrast to PC3 cells, the IC50 increases (efficacy decreases) with increasing PTX content for both DOTAP/DOPC/PTX and DLinTAP/DLinPC/PTX formulations. This effect is less pronounced for the DLinTAP/DLinPC/PTX formulations, and the efficacy of the DLinTAP/DLinPC/PTX formulation at 9 mol% PTX is lower than that of the formulation at 2 mol% PTX.

We again assessed DLinTAP/DLinPC lipid toxicity using a CL-only control. DLinTAP/DLinPC CLs without PTX caused >10% drop in cell viability only at or above lipid concentrations equivalent to those present when 75 nM PTX is delivered by CLs loaded with 2 mol% PTX (Figure 8A, blue line). In contrast, the IC50 for the same CLs with PTX is 17.6 nM PTX, about four times lower. It is therefore unlikely that lipid toxicity contributed significantly to the measured IC50 values for PTX-loaded CLs at any PTX content, given that the lipid/PTX ratio for the formulations decreased more than the IC50 increased (see below).

Viability of M21 cells as a function of delivered PTX concentration for the positive control, DOTAP/DOPC CLs loaded with 2 to 9 mol% PTX, is shown in Figure 8B. The decrease in cell viability with PTX concentration is very gradual and slower than that for PTX-loaded DLinTAP/DLinPC CLs (Figure 8A), suggesting that DLinTAP/DLinPC CLs induced cytotoxicity more effectively than DOTAP/DOPC CLs when used against M21 cells. According to literature data, DOTAP/DOPC CLs without PTX are not cytotoxic at concentrations well above those used in this experiment.^48^

For a more quantitative comparison, Figure 8C plots the IC50 for PTX cytotoxicity against M21 cells of the investigated formulations as a function of their membrane PTX content. The IC50 of PTX-loaded DLinTAP/DLinPC CLs was at least 2-fold lower than that of DOTAP/DOPC CLs at every PTX content tested. Because toxicity of PTX-free DLinTAP/DLinPC CLs is only observed well above the IC50 of those formulations, this improved efficacy is unlikely to be due to differences in lipid toxicity. Interestingly, the efficacy of formulations against M21 cells increases (their IC50 decreases) with PTX membrane solubility. This is in line with our expectations based on previous findings regarding the correlation of PTX solubility to cytotoxic efficacy.^48^

Taken together, PTX-loaded DLinTAP/DLinPC CLs show the same or better efficacy against cancer cells as the established DOTAP/DOPC/PTX formulation (Endo-TAG1), in addition to the improved drug loading capacity demonstrated by the kinetic phase diagrams.

## Conclusion

Our results have demonstrated how modifying lipid tail structure can improve cationic liposome (CL)-based carriers of cancer chemotherapy drug paclitaxel (PTX), by revealing two significant advantages over CL formulations that mimic the Endo-TAG1 (DOTAP/DOPC/PTX) vectors currently in phase III trials. The data show that replacement of lipid tails bearing one *cis* double bond (oleoyl tails, DO lipids) with tails bearing two *cis* double bonds (linoleoyl tails, DLin lipids) enhances solubility of PTX in CL vectors and impacts efficacy against in two human cancer cell lines. The increased tail unsaturation significantly improved PTX membrane solubility in the CLs from ≈3 mol% to ≈8 mol%. This improves the drug loading capacity of the CL vector, allowing the same amount of PTX to be delivered with much less lipid. Minimizing the amount of cationic lipids in *in vivo* delivery applications is highly desirable not only because it reduces cost but also because such cationic lipids can evoke a cellular immune response with measurable amounts of secreted interleukins.^109^

In addition to the important benefits derived from the reduced amount of cationic lipid needed to deliver PTX, a further significant finding of our study is that CL vectors based on DLin lipids have either similar or even improved efficacy when compared to vectors based on DO lipids, as evident by their IC50 of PTX cytotoxicity against prostate cancer (PC3) and melanoma (M21) cell lines, respectively. An increased efficacy allows reducing the amount of both lipid and PTX (or, alternatively, an enhanced effectiveness against cancer cells if the dose of PTX is kept constant).

Our results pave the path for *in vivo* studies to determine whether improved CL carrier properties, for PTX delivery to human cancer cells *in vitro*, translate to substantially improved outcomes *in vivo*. In particular, it will be important to establish whether the 8 to 10 hour solubility window, found for CLs with significantly larger PTX content, is sufficient for drug delivery *in vivo*.

We expect that this work will motivate future studies using chemical modifications of lipid structure as well as computational modeling to further explore how altering lipid tails affects their molecular affinity to PTX. This, when combined with kinetic phase diagrams as presented here, should lead to a comprehensive understanding of how lipid shape and tail structure correlates to PTX membrane solubility, paving the way to improved cancer therapeutics.

## Acknowledgement

This work was supported by the National Institutes of Health under Award R01GM130769 (mechanistic studies on the dependence of efficacy properties of cationic liposome vectors loaded with cancer drug paclitaxel on distribution of *cis* double bonds in lipid tails). Partial support was further provided by the US National Science Foundation (NSF) under Award DMR-1807327 (kinetic phase behavior of vesicles with hydrophobic molecules). VS was supported by the National Science Foundation Graduate Research Fellowship Program under Grant No. DGE 1144085. X-ray scattering work was carried out at the Stanford Synchrotron Radiation Lightsource, a Directorate of SLAC National Accelerator Laboratory and an Office of Science User Facility operated for the U.S. DOE Office of Science by Stanford University. We thank Rachel Behrens for help with the ESI-MS, which was performed at the Shared Experimental Facilities of the Materials Research Laboratory at UCSB. These facilities are supported by the MRSEC Program of the NSF under Award No. DMR 1720256; a member of the NSF-funded Materials Research Facilities Network (www.mrfn.org).

## Supplementary Material Contents

Detailed methods for the synthesis of DLinTAP; kinetic phase diagram and SAXS profile of DNA complexes of PTX-loaded CLs containing 70 mol% DOPE; tabulated cytotoxicity data; tables with the values for the IC50 and the slope factor of PTX cytotoxicity against PC3 and M21 cells.

## Data Availability

The raw/processed data required to reproduce these findings cannot be shared at this time as the data also forms part of an ongoing study.

