## Supplementary Material for "Enhanced Loading of Paclitaxel in Cationic Liposomes by Replacement of Oleoyl with Linoleoyl Lipid Tails with Benefits in Cancer Therapeutics from *In Vitro* Studies"

for

### Synthesis of DLinTAP

#### Materials and General Methods

Chemicals were purchased from Sigma-Aldrich at the highest available purity. Reactions were performed under an inert atmosphere of nitrogen. Silica gel from Fisher Scientific with a mesh size of 200-425 was used for flash chromatography.

For thin-layer chromatography (TLC), silica-covered aluminum sheets with fluorescence indicator (20 × 20 cm, Merck) were used. The TLC sheets were cut to the desired size using a razor blade. TLC plates were developed by heating after dipping into a solution of cerium ammonium molybdate (prepared by dissolving 5 g of ammonium pentamolybdate and 0.2 g of cerium(IV) sulfate in 100 mL of a mixture of sulfuric acid and water (1:9, v/v)).

Ultra-high pressure liquid chromatography–mass spectrometry (UPLC-MS) was performed using an Acquity UPLC H-Class chromatography system (Waters) coupled to a Xevo G2-X S ToF mass spectrometer (Waters) equipped with an ESI source and MassLynx software (version 4.1). The UPLC instrument consisted of a quaternary pump (Quaternary Solvent Manager), autosampler (Sample Manager-FTN) equipped with a 10 µL sample loop, and a tunable UV detector (TUV). Injections of 5 µL of the sample were separated through a Waters Acquity UPLC BEH C18 column (2.1 × 50 mm, 1.7 µm), under using isocratic elution with methanol (MS grade) at a flow rate of 0.4 mL/min. The column and the autosampler were maintained at 40°C and 10°C, respectively. ESI conditions were: source temperature 120°C, desolvation temperature 400°C, cone gas flow 20 L/h , desolvation gas flow 800 L/h , capillary voltage 1.5 kV, sampling cone 75, and source offset 80. The MS was acquired over an m/z range 50–2000 with a scan time equal to 0.5 s. Data were collected in the positive (ESI+) mode. Leucine Enkephalin ($[M + H]^+$ = 556.2771 m/z) (2 ng/µL) was used as lock mass for mass shift correction. The mass spectrometer was calibrated before analyses using a 2 µg/µL sodium iodide solution.

#### Synthesis Overview

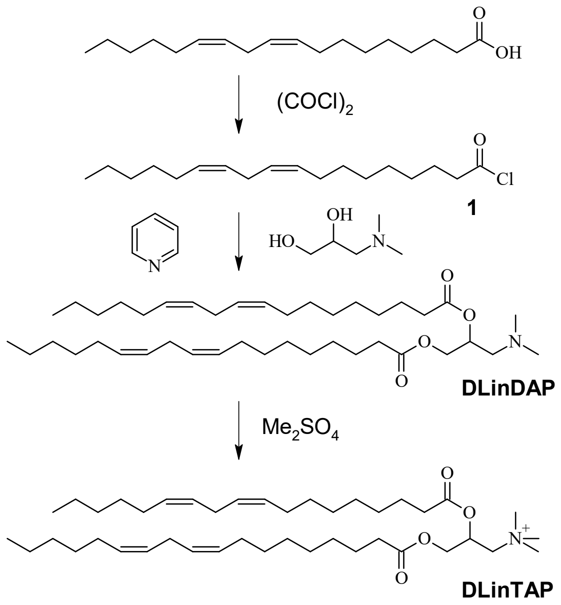

#### Procedures

**Linoleic acid chloride (1).**[1,2] To a solution of 2.57 g (9.16 mmol) linoleic acid in benzene (60 mL) was added 1 mL (11.7 mmol) of oxalyl chloride via syringe. The reaction mixture was stirred for 21 h at room temperature while monitoring gas evolution. Evaporation of volatiles in a vacuum yielded linoleic acid chloride as a colorless oil, which was used for the next step without further purification. $R_f$(hexanes/ethyl acetate=2:1, v/v)=0.95. (In the same eluent, $R_f$ of linoleic acid=0.59.)

**3-(dimethylamino)propane-1,2-diyl di((9*Z*,12*Z*)-octadeca-9,12-dienoate) (DLinDAP).** This step closely followed the procedure for preparation of 1,2-dioleoyloxy-3-(dimethylamino)propane by Leventis and Silvius.[3] To a solution of linoleic acid chloride (9.16 mmol, quantitative conversion assumed for previous step) and 431.6 mg (3.62 mmol) of 3-dimethylamino-1,2-propanediol in 40 mL of diethyl ether were added 339 µL (4.2 mmol) of dry pyridine. After stirring in the dark for 20 h, the reaction was quenched with methanol (1 mL) and volatiles were evaporated in a vacuum. The residue was dissolved in hexanes (75 mL), washed twice with 75 mL each of a 0.1 M solution of KOH in a mixture of methanol/water (1/1, v/v) and once with 0.1 M NaCl (63 mL). The hexane phase was separated, dried ($Na_2SO_4$), and evaporated in a vacuum. The residue was purified by flash chromatography on silica gel (130 g), eluting with hexanes/ethyl acetate (initially 4/1 (625 mL), then 3/1 (1000 mL) and 2/1 (300 mL); all v/v) to yield 1.663 g (2.582 mmol, 71%) of DLinDAP as a colorless oil. $R_f$(hexanes/ethyl acetate=2:1, v/v)=0.45.

**2,3-Dilinoleyloxy¬propyl¬trimethyl¬ammonium methylsufate (DLinTAP).** A solution of 1.434 g (2.227 mmol) DLinDAP and 1 mL (1.33 g, 10.5 mmol) of dimethylsulfate in acetone (10 mL) was stirred at 4 °C for 72 h. The resulting crystals were filtered off in the cold, briefly washed with cold acetone, and further purified by flash chromatography on silica gel (40 g). Elution with a gradient of methanol in chloroform (starting at 1:50, ending at 2:5, v/v) yielded 0.582 g (0.756 mmol, 34%) of DLinTAP as a colorless solid. $R_f$(chloroform/methanol/water=62:25:4, v/v/v): 0.62; $R_f$(chloroform/methanol=5:2, v/v): 0.52;[1] ESI-MS: 659.6162.

[1] The behavior of DLinTAP in TLC is somewhat capricious, with the $R_f$ showing a pronounced dependence on concentration and the not infrequent appearance of two spots (one of which, in some cases, was observed at very low $R_f$) for homogenous samples. A pure sample of commercial DOTAP exhibited similar behavior in TLC.

#### Cytotoxicity data

##### Normalized cell viability – PC3 cells

| | DOPC/DOTAP | | | | DLinPC/DLinTAP | | | | |
|---|---|---|---|---|---|---|---|---|---|
| **[PTX] /nM** | **2 mol% PTX** | **4 mol% PTX** | **6 mol% PTX** | **9 mol% PTX** | **Lipid Only** | **2 mol% PTX** | **4 mol% PTX** | **6 mol% PTX** | **9 mol% PTX** |
| **100** | 15.09 | 16.12 | 13.97 | 16.32 | 76.0 | 8.03 | 13.66 | 10.87 | 13.7 |
| **65** | 20.72 | 15.45 | 13.51 | 17.54 | 82.5 | 12.69 | 13.89 | 13.83 | 18.6 |
| **45** | 23.94 | 17.35 | 16.58 | 19.65 | 91.0 | 17.12 | 18.30 | 11.88 | 18.5 |
| **30** | 35.68 | 19.08 | 24.76 | 20.21 | 99.8 | 25.56 | 19.15 | 15.18 | 19.9 |
| **20** | 46.12 | 25.69 | 29.01 | 21.85 | 94.4 | 49.23 | 23.79 | 14.30 | 22.8 |
| **15** | 77.66 | 27.20 | 30.43 | 22.92 | 96.5 | 53.73 | 17.84 | 18.07 | 21.4 |
| **12.5** | 65.17 | 30.41 | 36.86 | 27.53 | 99.4 | 57.47 | 20.85 | 28.2 | 25.3 |
| **10** | 70.26 | 37.76 | 38.69 | 33.66 | 113.8 | 65.80 | 23.16 | 35.7 | 29.1 |
| **5** | 105.33 | 78.87 | 83.64 | 57.77 | | 109.47 | 59.09 | 71.0 | 57.4 |
| **1** | 117.99 | 85.54 | 105.87 | 93.93 | | 111.26 | 89.59 | 115.3 | 100.0 |

##### Standard error of normalized cell viability – PC3 cells

| | DOPC/DOTAP | | | | DLinPC/DLinTAP | | | | |
|---|---|---|---|---|---|---|---|---|---|
| **[PTX] /nM** | **2 mol% PTX** | **4 mol% PTX** | **6 mol% PTX** | **9 mol% PTX** | **Lipid Only** | **2 mol% PTX** | **4 mol% PTX** | **6 mol% PTX** | **9 mol% PTX** |
| **100** | 2.84 | 2.47 | 2.28 | 1.35 | 7.0 | 1.81 | 1.54 | 1.36 | 1.3 |
| **65** | 2.89 | 2.96 | 3.03 | 1.14 | 7.2 | 2.41 | 1.70 | 1.43 | 1.8 |
| **45** | 3.84 | 2.97 | 1.50 | 1.28 | 9.4 | 4.52 | 1.91 | 1.31 | 1.7 |
| **30** | 5.59 | 3.25 | 1.89 | 2.51 | 10.1 | 7.93 | 2.16 | 1.69 | 1.8 |
| **20** | 7.07 | 3.62 | 3.13 | 2.30 | 8.6 | 9.45 | 2.38 | 2.04 | 2.1 |
| **15** | 11.37 | 3.96 | 3.10 | 4.72 | 9.2 | 16.37 | 3.87 | 2.86 | 2.0 |
| **12.5** | 10.37 | 5.70 | 2.43 | 4.51 | 9.1 | 13.62 | 4.21 | 2.8 | 2.2 |
| **10** | 10.14 | 5.38 | 2.60 | 7.51 | 13.6 | 14.24 | 3.65 | 3.3 | 3.9 |
| **5** | 14.84 | 13.82 | 6.27 | 16.27 | | 12.79 | 6.23 | 6.3 | 8.1 |
| **1** | 17.56 | 13.38 | 5.11 | 23.82 | | 12.01 | 9.59 | 11.1 | 10.6 |

#### Normalized cell viability – M21 cells

| | DOPC/DOTAP | | | | DLinPC/DLinTAP | | | | |
|---|---|---|---|---|---|---|---|---|---|
| **[PTX] /nM** | **2 mol% PTX** | **4 mol% PTX** | **6 mol% PTX** | **9 mol% PTX** | **Lipid Only** | **2 mol% PTX** | **4 mol% PTX** | **6 mol% PTX** | **9 mol% PTX** |
| **500** | 15.73 | 23.97 | 17.63 | 21.47 | 16.2 | 9.89 | 12.55 | 12.60 | 14.5 |
| **100** | 26.31 | 64.49 | 60.25 | 61.80 | 54.9 | 15.12 | 24.16 | 26.44 | 36.0 |
| **75** | 36.57 | 76.66 | 58.66 | 64.93 | 82.9 | 18.76 | 30.79 | 30.70 | 44.5 |
| **65** | 38.61 | 82.99 | 70.49 | 67.72 | 97.3 | 25.94 | 44.60 | 36.28 | 42.5 |
| **55** | 60.09 | 72.50 | 63.23 | 53.31 | 93.9 | 25.16 | 49.15 | 40.49 | 50.5 |
| **45** | 60.06 | 96.69 | 77.49 | 65.07 | 92.0 | 27.80 | 46.73 | 52.39 | 55.8 |
| **30** | 85.75 | 98.04 | 81.05 | 67.01 | 95.3 | 30.40 | 54.35 | 56.9 | 60.8 |
| **20** | 101.24 | 103.32 | 88.68 | 66.02 | 96.3 | 83.27 | 63.45 | 77.6 | 78.0 |
| **10** | 95.64 | 95.10 | 102.14 | 88.24 | | 85.54 | 70.34 | 93.8 | 111.7 |
| **3** | 129.22 | 152.66 | 117.74 | 98.49 | | 110.32 | 121.21 | 128.2 | 129.5 |

#### Standard error of normalized cell viability – M21 cells

| | DOPC/DOTAP | | | | DLinPC/DLinTAP | | | | |
|---|---|---|---|---|---|---|---|---|---|
| **[PTX] /nM** | **2 mol% PTX** | **4 mol% PTX** | **6 mol% PTX** | **9 mol% PTX** | **Lipid Only** | **2 mol% PTX** | **4 mol% PTX** | **6 mol% PTX** | **9 mol% PTX** |
| **500** | 1.57 | 2.29 | 1.6 | 1.63 | 4.60 | 0.66 | 0.79 | 1.26 | 1.17 |
| **100** | 1.44 | 4.26 | 4.6 | 9.79 | 2.90 | 1.81 | 1.64 | 1.85 | 3.00 |
| **75** | 3.80 | 7.35 | 4.57 | 5.14 | 4.18 | 3.02 | 2.68 | 2.85 | 4.60 |
| **65** | 5.08 | 9.19 | 9.41 | 5.53 | 2.38 | 2.47 | 4.29 | 1.48 | 2.90 |
| **55** | 7.24 | 13.02 | 6.43 | 2.43 | 3.53 | 1.22 | 4.52 | 3.89 | 4.18 |
| **45** | 12.83 | 7.22 | 10.05 | 3.52 | 3.25 | 1.12 | 1.48 | 4.44 | 2.38 |
| **30** | 16.38 | 6.33 | 9.35 | 7.14 | 5.60 | 3.85 | 2.97 | 5.82 | 3.53 |
| **20** | 12.94 | 3.81 | 9.01 | 4.33 | 12.90 | 9.43 | 3.56 | 6.81 | 3.25 |
| **10** | 8.22 | 3.12 | 10.25 | 9.76 | | 6.98 | 2.96 | 3.69 | 5.60 |
| **3** | 5.86 | 9.26 | 7.78 | 10.33 | | 6.97 | 6.92 | 6.64 | 12.90 |

#### Kinetic Phase Diagram for PTX-loaded DOTAP/DOPE CLs

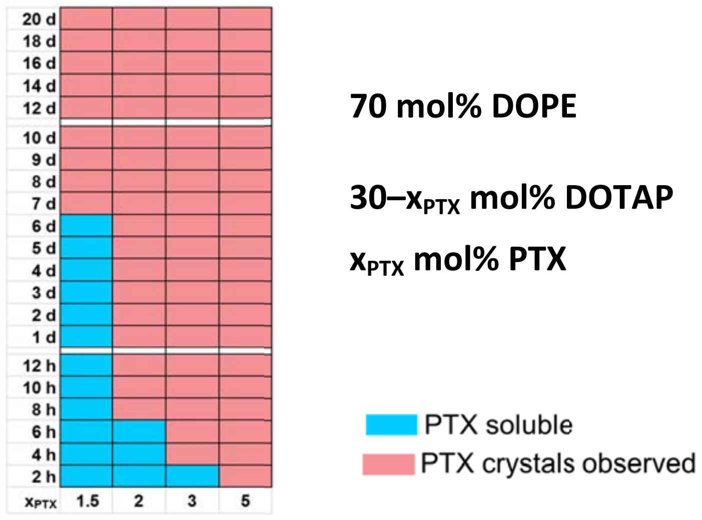

**Figure S1.** Kinetic phase diagram of PTX solubility in PTX-loaded unsonicated CLs of molar composition DOTAP/DOPE/PTX=30–$x_{PTX}$:70:$x_{PTX}$. Differential-interference-contrast microscopy was used to detect PTX crystallization at the indicated times after hydration. The blue color indicates absence of PTX crystals (i.e., PTX remained soluble in the membranes), while the red color indicates presence of PTX crystals. The PTX membrane solubility boundary (the border between the blue and red crystallized areas) was determined from the median of 3 to 5 separate trials at each PTX content. The DNA complexes of membranes with molar composition DOTAP/DOPE/PTX=28:70:2 form the inverse hexagonal ($H_{II}^{C}$) phase (see Figure S2). The low PTX solubility in these membranes demonstrates that it is not the self-assembled structure of the lipid that increases the PTX solubility in DLinTAP/DLinPC CLs.

#### SAXS profiles for PTX-loaded DOTAP/DOPC and DOTAP/DOPE CLs

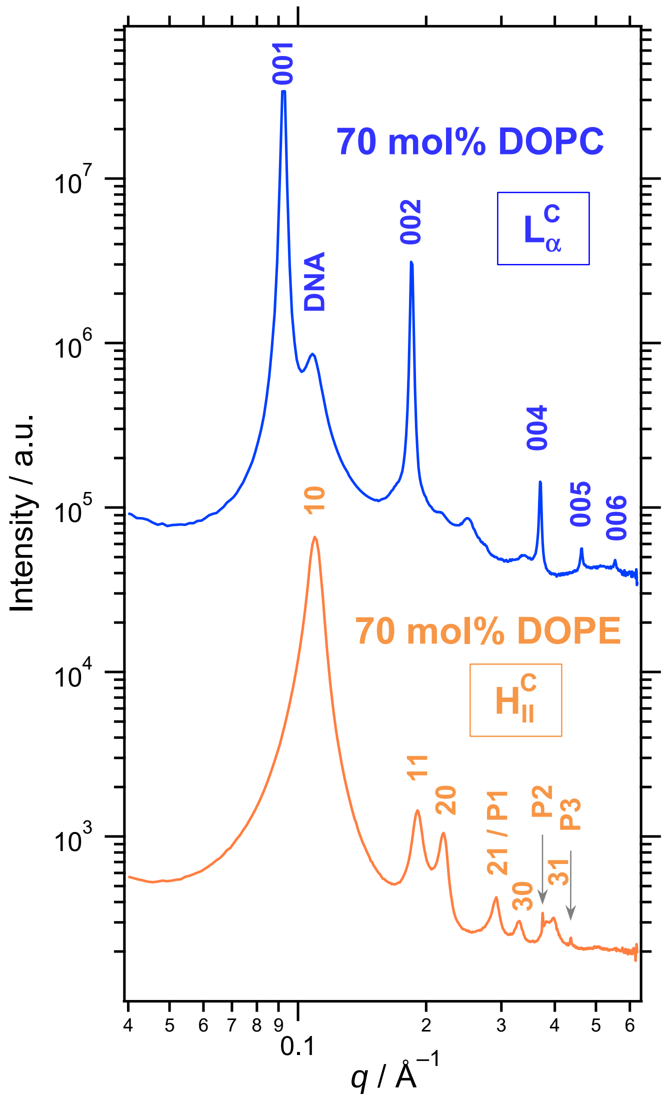

**Figure S2.** Small-angle X-ray scattering profiles of DNA complexes of PTX-loaded CLs of molar compositions DOTAP/DOPC/PTX= 70/28/2 (blue, top) and DOTAP/DOPE/PTX=70/28/2 (orange, bottom), revealing the self-assembled structures. Peak assignments are shown on the plots as follows: $L_\alpha^C$ (00L: 001, 002, 003, 004, 005, 006), $H_{II}^C$ (HK: 10, 11, 20, 21, 30, 31), DNA–DNA interaxial spacing in the $L_\alpha^C$ phase (DNA) and crystallized PTX (P1, P2, P3). CLs were complexed with calf thymus DNA at a 1:1 charge ratio.

#### Values of IC50 and Slope Factor

**Table S1.** The IC50 of PTX cytotoxicity against PC3 cells for PTX-loaded CLs with differing tails structure and PTX membrane content. The values of IC50 and the slope factor were determined from the plots of (normalized) cell viability against PTX concentration (Figure 7) as described in the Methods section.

| | IC50 | | Slope Factor | |
|---|---|---|---|---|
| $x_{PTX}$ | DLin CLs | DO CLs | DLin CLs | DO CLs |
| 2 | 13.4 nM | 13.2 nM | 2.01 | 1.88 |
| 4 | 4.8 nM | 7.7 nM | 2.16 | 2.47 |
| 6 | 5.9 nM | 6.8 nM | 2.45 | 1.79 |
| 9 | 4.0 nM | 5.1 nM | 1.48 | 1.94 |

**Table S2.** The IC50 of PTX cytotoxicity against M21 cells for PTX-loaded CLs with differing tails structure and PTX membrane content. The values of IC50 and the slope factor were determined from the plots of (normalized) cell viability against PTX concentration (Figure 8) as described in the Methods section.

| | IC50 | | Slope Factor | |
|---|---|---|---|---|
| $x_{PTX}$ | DLin CLs | DO CLs | DLin CLs | DO CLs |
| 2 | 17.6 nM | 35.5 nM | 1.62 | 2.01 |
| 4 | 19.7 nM | 39.1 nM | 1.15 | 1.12 |
| 6 | 23.1 nM | 56.3 nM | 1.51 | 1.48 |
| 9 | 27.7 nM | 54.8 nM | 1.29 | 1.18 |
